## Supplementary materials for "Role of individual and population heterogeneity in shaping dynamics of multi-pathogen shedding in an island endemic bat"

**S1 Table. Summary of the statistical models (models M1 to M4) used to analyse single shedding dynamics in *M. francoismoutoui.*** Significant variables are in bold and the asterisk represents the interaction between two variables. All GAMs were fitted with a binomial distribution. Model M4 (GLM) was fitted with a Gaussian distribution, and its performance was compared to a null model using AIC criterion. The percentage of deviance explained was calculated by comparing full model with null model. PMV: Paramyxovirus, LEPTO: *Leptospira* bacteria, HSV: Herpesvirus.

| *Type and model number* | *Levels and number of individuals* | *Response variable* | *Deviance explained (%)* | *Explanatory variables* | *EDF* | *Chi²* | *Estimate (± SE)* | *Odds ratios* | *Z value* | *P* |
|---|---|---|---|---|---|---|---|---|---|---|
| GAM M1 | All individuals N = 5518 | PMV | 14.1 | **s(Sampling period)** | 5.70 | 15.05 | | | | 0.032 |
| | | | | s(SSAS) | 4.32 | 1.55 | | | | 0.90 |
| | | | | logSize | | | 0.01 (±0.03) | 1.01 | 0.37 | 0.71 |
| | | | | **Age** | | | -3.46 (±0.59) | 0.03 | -5.83 | $5.62^{-09}$ |
| | | | | **Sex** | | | -0.45 (±0.09) | 0.64 | -5.03 | $4.81^{-07}$ |
| | | | | Age*Sex | | | 0.99 (±0.75) | 2.70 | 1.33 | 0.18 |
| | | | | **LEPTO** | | | 2.54 (±0.64) | 12.62 | 3.96 | $7.47^{-05}$ |
| | | | | **HSV** | | | 2.65 (±0.52) | 14.13 | 5.12 | $2.99^{-07}$ |
| | | | | **LEPTO*HSV** | | | -2.00 (±0.65) | 0.14 | -3.10 | 0.002 |
| GAM M2 | All individuals N = 5518 | LEPTO | 10.3 | s(Sampling period) | 6.17 | 12.66 | | | | 0.09 |
| | | | | s(SSAS) | 2.89 | 1.76 | | | | 0.60 |
| | | | | logSize | | | 0.02 (±0.03) | 1.02 | 0.80 | 0.43 |
| | | | | **Age** | | | -1.41 (±0.23) | 0.24 | -6.17 | $6.73^{-10}$ |
| | | | | Sex | | | -0.07 (±0.09) | 0.94 | -0.77 | 0.44 |
| | | | | Age*Sex | | | 0.04 (±0.29) | 1.04 | 0.14 | 0.89 |
| | | | | **PMV** | | | 2.09 (±0.62) | 8.10 | 3.35 | $7.99^{-04}$ |
| | | | | **HSV** | | | 1.12 (±0.19) | 3.05 | 5.89 | $3.95^{-09}$ |

| | | | | | | | | | |
|---|---|---|---|---|---|---|---|---|---|
| | | | | **PMV*HSV** | | | -1.54 (±0.62) | 0.21 | -2.46 | 0.01 |
| GAM<br>M3 | All individuals<br>N = 3981 | HSV | 44.9 | **s(Sampling period)**<br>s(SSAS)<br>**logSize**<br>**Age**<br>**Sex**<br>**Age*Sex**<br>**PMV**<br>**LEPTO**<br>**PMV*LEPTO** | 6.67<br>4.93 | 19.86<br>7.38 | <br><br>-0.14 (±0.05)<br>-2.80 (±0.23)<br>0.81 (±0.21)<br>-0.88 (±0.29)<br>2.52 (±0.52)<br>1.09 (±0.19)<br>1.59 (±0.63) | <br><br>0.87<br>0.06<br>2.25<br>0.41<br>12.47<br>2.98<br>0.20 | <br><br>-2.83<br>11.88<br>3.84<br>-3.03<br>4.84<br>5.63<br>-2.51 | 0.01<br>0.40<br>0.005<br>$2^{-16}$<br>$1.25^{-04}$<br>0.002<br>$1.30^{-06}$<br>$1.81^{-08}$<br>0.01 |
| GAM<br>M3bis | Adults only<br>N = 3344 | HSV | 14.8 | s(Sampling period)<br>s(SSAS)<br>logSize<br>**Sex**<br>**PMV**<br>**LEPTO**<br>**PMV*LEPTO** | 5.90<br>4.59 | 11.37<br>1.57 | <br><br>-0.08 (±0.09)<br>0.96 (±0.22)<br>2.30 (±0.52)<br>1.38 (±0.28)<br>-1.74 (±0.66) | <br><br>0.93<br>2.62<br>9.93<br>3.98<br>0.18 | <br><br>-0.83<br>4.44<br>4.42<br>4.87<br>-2.63 | 0.14<br>0.93<br>0.41<br>$9.06^{-06}$<br>$9.75^{-06}$<br>$1.12^{-06}$<br>0.009 |
| GLM<br>M4 | *Leptospira* PCR-positive individuals<br>N = 2529 | Ct LEPTO | 5.2 | **Age (AIC = 14036)**<br>Null (AIC = 14169) | | | 4.53 (±0.38) | | 11.78 | $2^{-16}$ |

**S2 Table. Summary of the statistical models (models M5 to M12) used to analyse the mono-excretion dynamics in *M. francoismoutoui* during pregnancy and mating periods, specifically.** GLMs were fitted with a binomial distribution, except for Ct LEPTO variable fitted with gaussian distribution. Final models (in bold) were selected by comparing full model with null model and using best AIC criterion (when ΔAIC>2). The percentage of deviance explained was calculated by comparing full model with null model. PMV: Paramyxovirus, LEPTO: *Leptospira* bacteria, HSV: Herpesvirus. M0: female with no visible nipples.

| *Type and model number* | *Levels and number of individuals* | *Deviance explained (%)* | *Response variable* | *Explanatory variables (AIC)* | *Estimate (± SE)* | *Z value* | *P* |
|---|---|---|---|---|---|---|---|
| GLM M5 | Adult females N = 523 | 5.2 | PMV | **Pregnancy (684)** Null (720) | 1.49 (±0.26) | 5.76 | $8.31^{-09}$ |
| GLM M5bis | Adult females, without non-pregnant M0 N = 470 | 0.1 | PMV | Pregnancy (630) **Null (628)** | 0.25 (±0.35) | 0.72 | 0.47 |
| GLM M6 | Adult female N = 522 | 4.1 | LEPTO | **Pregnancy (682)** Null (709) | 1.27 (±0.25) | 5.20 | $2.02^{-07}$ |
| GLM M6bis | Adult females, without non-pregnant M0 N = 469 | 0.3 | LEPTO | Pregnancy (618) **Null (618)** | 0.47 (±0.35) | 1.35 | 0.18 |
| GLM M7 | Adult females N = 307 | 0.9 | Ct LEPTO | Pregnancy (1713) **Null (1713)** | -1.27 (±0.75) | -1.69 | 0.09 |
| GLM M7bis | Adult females, without non-pregnant M0 N = 296 | 0.01 | Ct LEPTO | Pregnancy (1637) **Null (1635)** | 0.16 (±0.91) | 0.18 | 0.86 |
| GLM | Adult females | 1.4 | HSV | Pregnancy (102) | 0.78 (±0.64) | 1.22 | 0.22 |

| | | | | | | | |
|---|---|---|---|---|---|---|---|
| M8 | N = 377 | | | **Null (101)** | | | |
| GLM<br>M8bis | Adult females, without non-pregnant M0<br>N = 331 | 2.2 | HSV | Pregnancy (70)<br>**Null (70)** | -15.84 (±1872) | -0.008 | 0.99 |
| GLM<br>M9 | Adult males<br>N = 353 | 0.001 | PMV | Reproduction (485)<br>**Null (483)** | 0.02 (±0.22) | 0.09 | 0.93 |
| GLM<br>M10 | Adult males<br>N = 353 | 1.5 | LEPTO | **Reproduction (483)**<br>Null (489) | 0.59 (±0.22) | 2.70 | 0.007 |
| GLM<br>M11 | Adult males<br>N = 163 | 1.6 | Ct LEPTO | Reproduction (898)<br>**Null (899)** | 0.96 (±0.59) | 1.62 | 0.11 |
| GLM<br>M12 | Adult males<br>N = 228 | 4.4 | HSV | Reproduction (63)<br>**Null (64)** | 1.55 (±1.09) | 1.42 | 0.16 |

**S3 Table. Summary of the statistical models (models M13 to M16) used to analyse dual and triple shedding dynamics in *M. francoismoutoui*.** Significant variables are in bold and the asterisk represents the interaction between two variables. The percentage of deviance explained was calculated by comparing full model with null model. All GAMs were fitted with a binomial distribution. PMV: Paramyxovirus, LEPTO: *Leptospira* bacteria, HSV: Herpesvirus.

| *Type and model number* | *Levels and number of individuals* | *Response variable* | *Deviance explained (%)* | *Explanatory variables* | *EDF* | *Chi²* | *Estimate (± SE)* | *Z value* | *P* |
|---|---|---|---|---|---|---|---|---|---|
| GAM M13 | Individuals tested for the three infectious agents N = 3784 | PMV - LEPTO | 9.1 | s(Sampling period) | 5.45 | 5.68 | | | 0.52 |
| | | | | s(SSAS) | 4.17 | 3.71 | | | 0.67 |
| | | | | logSize | | | 0.04 (±0.04) | 1.02 | 0.31 |
| | | | | **Age** | | | -4.75 (±1.01) | -4.73 | $2.30^{-06}$ |
| | | | | **Sex** | | | -0.49 (±0.10) | -5.04 | $4.70^{-07}$ |
| | | | | Age*Sex | | | 0.46 (±1.42) | 0.33 | 0.75 |
| GAM M14 | Individuals tested for the three infectious agents N = 3784 | LEPTO - HSV | 10.8 | s(Sampling period) | 5.81 | 11.67 | | | 0.14 |
| | | | | s(SSAS) | 4.39 | 2.91 | | | 0.71 |
| | | | | logSize | | | 0.02 (±0.03) | 0.52 | 0.61 |
| | | | | **Age** | | | -2.81 (±0.26) | -10.67 | $2^{-16}$ |
| | | | | Sex | | | -0.06 (±0.08) | -0.67 | 0.51 |
| | | | | Age*Sex | | | -0.26 (±0.39) | -0.67 | 0.50 |
| GAM M15 | Individuals tested for the three infectious agents N = 3784 | PMV -HSV | 11.0 | **s(Sampling period)** | 5.74 | 17.46 | | | 0.01 |
| | | | | s(SSAS) | 4.34 | 1.94 | | | 0.86 |
| | | | | logSize | | | 0.007 (±0.03) | 0.22 | 0.83 |
| | | | | **Age** | | | -4.33 (±0.59) | -7.39 | $1.43^{-13}$ |
| | | | | **Sex** | | | -0.37 (±0.09) | -4.26 | $2.01^{-05}$ |
| | | | | Age*Sex | | | 0.88 (±0.74) | 1.19 | 0.24 |
| GAM M16 | Individuals tested for the three infectious agents | PMV – LEPTO - HSV | 8.9 | s(Sampling period) | 5.44 | 5.33 | | | 0.56 |
| | | | | s(SSAS) | 4.17 | 4.21 | | | 0.62 |
| | | | | logSize | | | 0.04 (±0.04) | 1.00 | 0.32 |

| | | | | | | | | | |
|---|---|---|---|---|---|---|---|---|---|
| | N = 3784 | | | **Age**<br>**Sex**<br>Age*Sex | | | -4.72 (±1.01)<br>-0.47 (±0.10)<br>0.44 (±1.42) | -4.69<br>-4.76<br>0.31 | $2.68^{-06}$<br>$1.98^{-06}$<br>0.76 |

**S4 Table. Summary of the statistical models (models M17 to M24) used to analyse dual and triple shedding dynamics in *M. francoismoutoui* during pregnancy and mating periods specifically.** GLMs were fitted with a binomial distribution. Final models (in bold) were selected by comparing full model with null model and using best AIC criterion (when ΔAIC>2). The percentage of deviance explained was calculated by comparing full model with null model. PMV: Paramyxovirus, LEPTO: *Leptospira* bacteria, HSV: Herpesvirus. M0: female with no visible nipples.

| *Type and model number* | *Levels and number of individuals* | *Response variable* | *Deviance explained (%)* | *Explanatory variables (AIC)* | *Estimate (± SE)* | *Z value* | *P* |
|---|---|---|---|---|---|---|---|
| GLM M17 | Adult females<br>N = 370 | PMV - LEPTO | 4.3 | **Pregnancy (456)**<br>Null (474) | 1.39 (±0.35) | 4.01 | $6.15^{-05}$ |
| GLM M17bis | Adult females, without non-pregnant M0<br>N = 323 | PMV - LEPTO | 0.2 | Pregnancy (432)<br>**Null (431)** | 0.38 (±0.40) | 0.97 | 0.33 |
| GLM M18 | Adult females<br>N = 370 | LEPTO - HSV | 5.2 | **Pregnancy (480)**<br>Null (504) | 1.32 (±0.27) | 4.93 | $8.05^{-07}$ |
| GLM M18bis | Adult females, without non-pregnant M0<br>N = 323 | LEPTO - HSV | 0.4 | Pregnancy (423)<br>**Null (423)** | 0.46 (±0.37) | 1.24 | 0.22 |
| GLM M19 | Adult females<br>N = 370 | PMV - HSV | 5.6 | **Pregnancy (487)**<br>Null (514) | 1.42 (±0.28) | 5.07 | $3.97^{-07}$ |
| GLM M19bis | Adult females without non-pregnant M0<br>N = 323 | PMV - HSV | 0.1 | Pregnancy (441)<br>**Null (439)** | 0.21 (±0.37) | 0.57 | 0.57 |
| GLM M20 | Adult females<br>N = 370 | PMV - LEPTO - HSV | 3.8 | **Pregnancy (451)**<br>Null (467) | 1.31 (±0.35) | 3.79 | $1.5^{-04}$ |

| | | | | | | | |
|---|---|---|---|---|---|---|---|
| GLM M20bis | Adult females, without non-pregnant M0 N = 323 | PMV - LEPTO - HSV | 0.1 | Pregnancy (427) **Null (426)** | 0.31 (±0.40) | 0.78 | 0.43 |
| GLM M21 | Adult males N = 214 | PMV - LEPTO | 0.3 | Reproduction (231) **Null (230)** | 0.29 (±0.33) | 0.89 | 0.38 |
| GLM M22 | Adult males N = 214 | LEPTO - HSV | 2.4 | **Reproduction (293)** Null (298) | 0.75 (±0.28) | 2.66 | 0.008 |
| GLM M23 | Adult males N = 214 | PMV -HSV | 0.1 | Reproduction (297) **Null (295)** | -0.18 (±0.28) | -0.63 | 0.53 |
| GLM M24 | Adult males N = 214 | PMV – LEPTO - HSV | 0.3 | Reproduction (231) **Null (230)** | 0.29 (±0.33) | 0.89 | 0.38 |

**S5 Table. Summary of the statistical models (models M25 and M26) used to analyze recapture data in *M. francoismoutoui*.** Transitions in Herpesvirus shedding status was not analysed because not enough negative saliva samples were available. Interval: time interval (days) between recaptures. Repro: reproductive status transition (active to active, non-active to active, active to non-active and non-active to non-active). PMV: Paramyxovirus, LEPTO: *Leptospira* bacteria.

| *Type and model number* | *Levels and number of individuals* | *Response variable* | *Explanatory variables* | *AIC* |
|---|---|---|---|---|
| Multinomial<br>M25 (a-h) | All individuals recaptured at least one time<br>N = 457 | LEPTO | a: Interval + Sex + Repro<br>b: Sex + Repro<br>c: Interval + Sex<br>d: **Interval + Repro**<br>e: Sex<br>f: Repro<br>g: Interval<br>h: Null | 1145<br>1157<br>1144<br>1140<br>1155<br>1152<br>1142<br>1153 |
| Multinomial<br>M26 (a-h) | All individuals recaptured at least one time<br>N = 471 | PMV | a: Interval + Sex + Repro<br>b: Sex + Repro<br>c: Interval + Sex<br>d: Interval + Repro<br>e: Sex<br>f: **Repro**<br>g: Interval<br>h: Null | 1123<br>1122<br>1130<br>1119<br>1128<br>1118<br>1134<br>1132 |

**S6 Table. Characteristics and coding of sampling periods used in GAMs.**

| Year | Sampling period | Sampling dates (MM/DD) | GAM code |
|---|---|---|---|
| 2019 | January - February | From 01/24 to 02/27 | 1 |
| | March - April | From 03/07 to 04/09 | 2 |
| | April - May | From 04/23 to 05/13 | 3 |
| | June | From 06/04 to 06/19 | 4 |
| | July | From 07/16 to 07/30 | 5 |
| | September | From 09/03 to 09/17 | 6 |
| | October - November | From 10/21 to 11/07 | 7 |
| | November - December | From 11/25 to 12/10 | 8 |
| 2020 | January - February | From 01/15 to 02/05 | 9 |
| | March | From 03/02 to 03/20 | 10 |
| | May - June | From 05/19 to 06/04 | 11.5 |
| | July | From 07/06 to 07/21 | 13 |
| | September | From 09/07 to 09/24 | 14 |
| | October | From 10/14 to 10/29 | 15 |
| | November - December | From 11/23 to 12/09 | 16 |

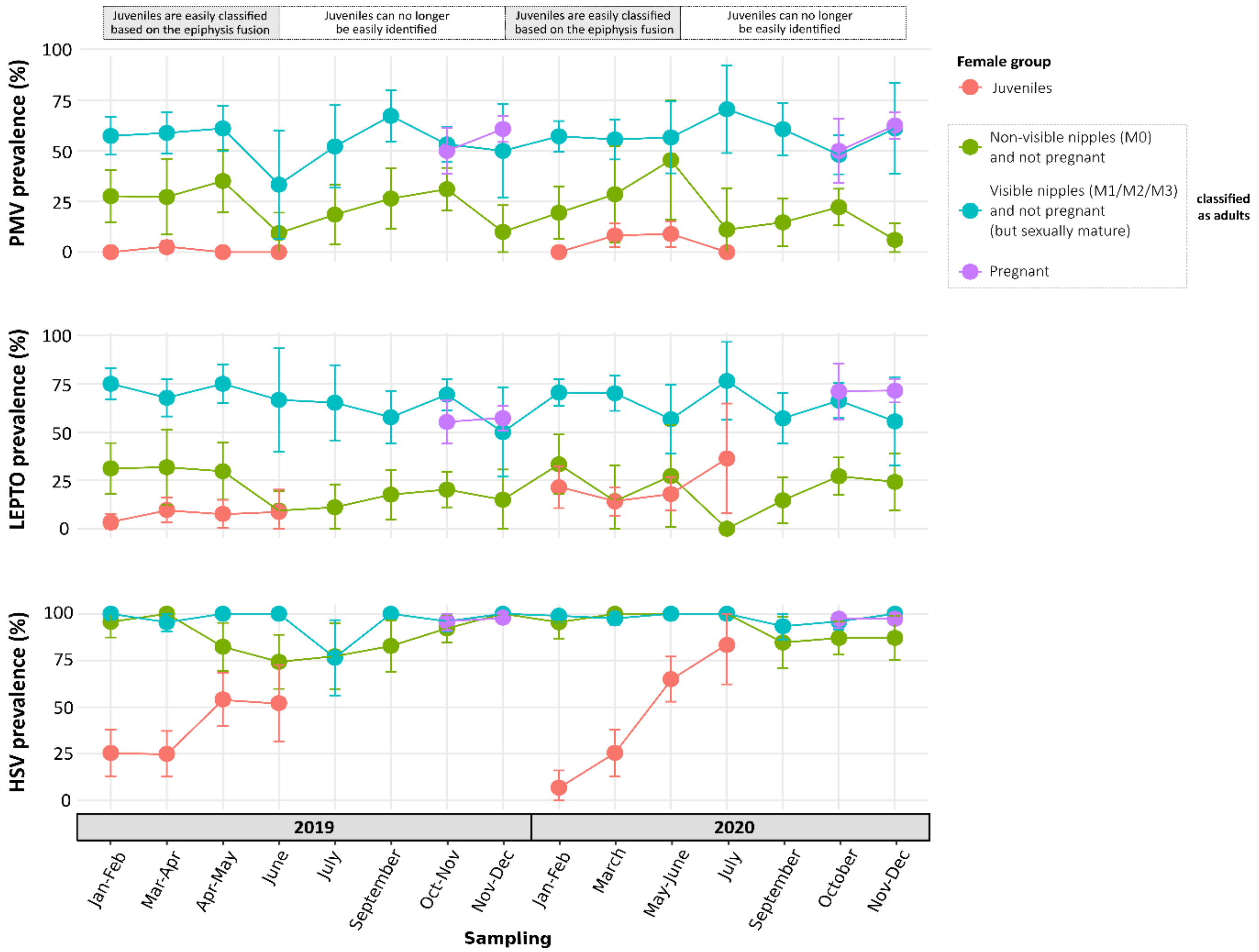


**S1 Fig. Temporal variation of paramyxovirus (PMV), *Leptospira* (LEPTO) and Herpesvirus (HSV) prevalence in *M. francoismoutoui* females, according to age and reproductive characteristics (pregnancy and visibility of nipples).** Note that in the green group, bats can include both adult bats that were not sexually mature yet, as well as misclassified juveniles.

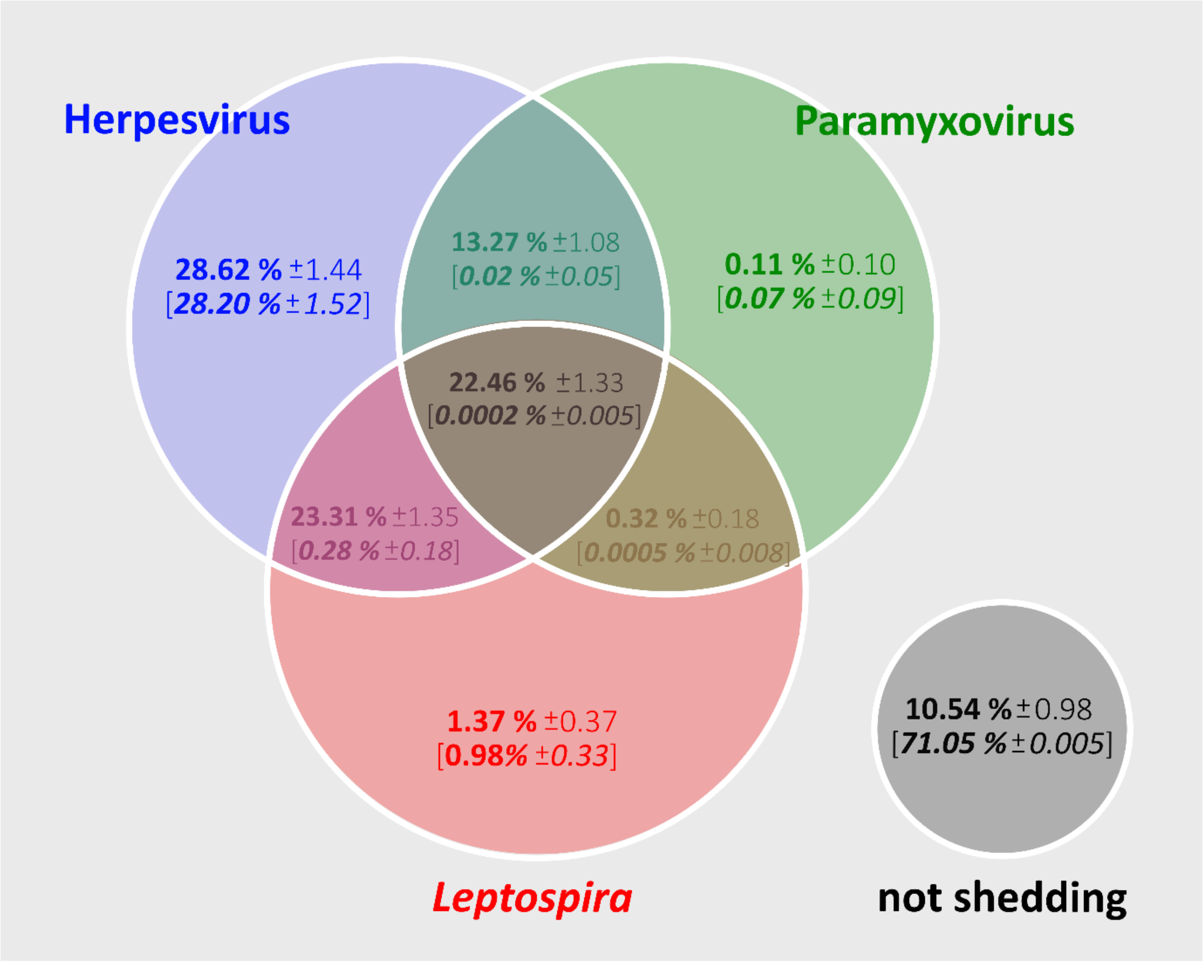


**S2 Fig. Venn diagram of *M. francoismoutoui* bats tested for the three infectious agents (n = 3784).** Observed proportions are shown with 95% confidence intervals (CI), and expected prevalence and 95% CI are indicated below in square brackets.

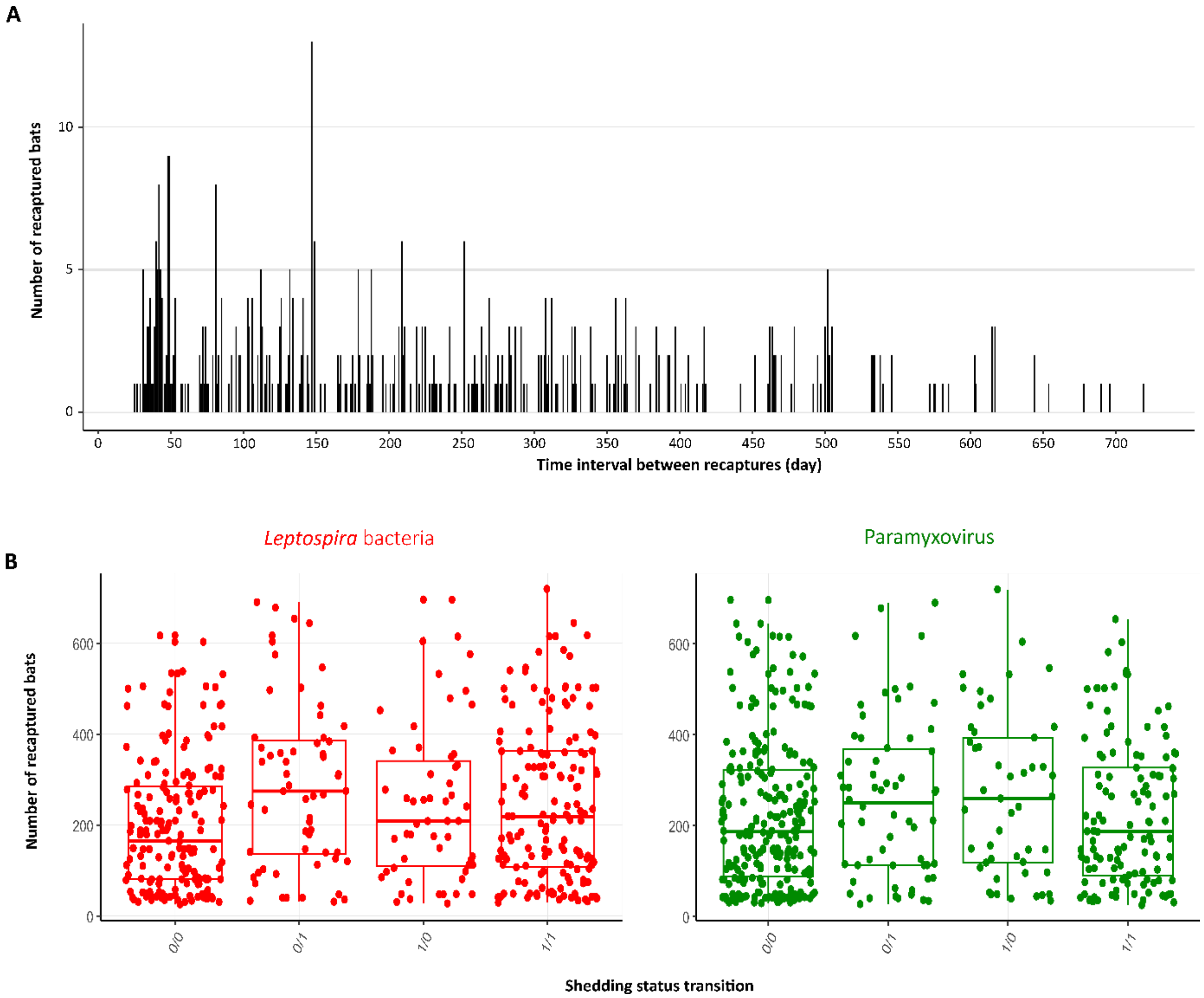


**S3 Fig. Details of recaptured *M. francoismoutoui* bats.** (A) Distribution of time interval between recaptures. (B) Variation of time intervals across the four categories of shedding status transitions, for *Leptospira* and paramyxovirus. Shedding status is coded with 0 for non-shedding and 1 for shedding bats.

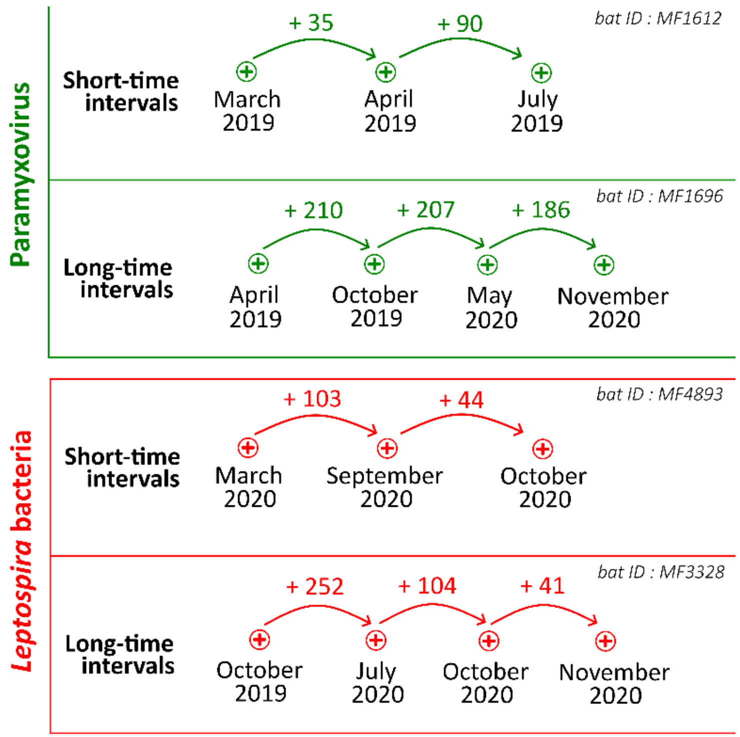


**S4 Fig. Examples of recaptured bats with always-positive status for paramyxovirus and *Leptospira*.** The time interval (in days) between two recaptures is indicated above the arrow.
